# Single-cell imaging links antibiotic resistance gene copy number and bacterial host identity

**DOI:** 10.64898/2026.11.10.765748

**Authors:** Gangan Wang, Liliana Cristina Moraru, Ismail Hayani Aji, Florin Musat, Jochen A. Müller, Niculina Musat

## Abstract

Identifying the *in situ* microbial hosts of antibiotic resistance genes (ARGs) in complex environments remains a major challenge, as many of these genes are associated with mobile genetic elements. Here, we introduce combined-FISH, which integrates direct-geneFISH for ARG detection with catalyzed reporter deposition fluorescence *in situ* hybridization (CARD-FISH) for rRNA labelling, thereby enabling the simultaneous visualization and copy number quantification of ARGs in taxonomically identified single cells within microbial samples. Using the sulfonamide resistance gene *sul*1 as a model target, we designed a polynucleotide probe set covering all non-redundant database sequences of this gene within 5% sequence mismatch. The method was validated using model organisms harboring plasmids with *sul*1, *Escherichia coli* K12 J53 with R388 and *Acinetobacter defluvii* with pOXA58_010030. Combined-FISH enabled quantitative determination of *sul*1 copy numbers at single-cell resolution, with *E. coli* harboring 1–8 copies and *A. defluvii* 1–4 copies per cell. Following plasmid-mediated transfer of *sul*1 from *E. coli* to *Pseudomonas alloputida*, recipient cells typically contained 2–6 *sul*1 copies, with up to 14 copies detected in individual cells, demonstrating the applicability of the approach for investigating horizontal gene transfer. The method was successfully applied to river biofilm samples containing *sul*1-harboring bacteria, enabling simultaneous tracking of the bacterial host and its associated ARG within a complex microbial community. Collectively, these results establish combined-FISH as a powerful tool for quantitatively linking ARGs and microbial hosts *in situ*, and for investigating plasmid copy number and gene transfer at single-cell resolution in technical and environmental systems.

## Introduction

Antimicrobial resistance (AMR) has been recognized by the World Health Organization as one of the most serious global public health threats of the 21^st^ century ^1^. To curtail the global spread of AMR, the One Health approach advocates for the inclusion of environmental compartments in surveillance and prevention efforts ^2^. However, effective risk assessment requires not only detecting the presence of antibiotic resistance genes (ARGs) but also identifying their microbial hosts and mobility potential ^3^. Linking ARGs and host identity is therefore essential for understanding the dissemination of resistance in natural environments and can further support the development of targeted strategies to mitigate the environmental spread of AMR.

Numerous approaches have been developed to investigate ARG-host associations in environmental microbiomes, each with distinct strengths and limitations. Culture-based methods provide direct validation of resistance phenotypes and their genetic determinants but are restricted by the low cultivability of environmental microorganisms and are often labor-intensive ^4–6^. Culture-independent sequencing approaches, particularly metagenomics, provide comprehensive insights into ARG diversity and abundance. However, as ARGs are frequently associated with mobile genetic elements (MGEs) such as plasmids, it remains a challenge to link resistance genes to their microbial hosts using conventional metagenomic approaches alone ^3,7^. Host assignment can be improved by methods based on intracellular DNA cross-linking prior to extraction and sequencing, such as Hi-C, which preserve physical associations between ARGs and host genomes. Nevertheless, these approaches remain susceptible to assembly and linkage biases, are cost-intensive, and generally lack true single-cell resolution ^8–10^.

Single-cell approaches have partially addressed these limitations. For example, epicPCR (emulsion, paired isolation, and concatenation PCR) enables high-throughput linkage of ARGs with 16S rRNA genes through the generation and parallel sequencing of fusion amplicons ^3,11–13^. However, the method provides only semi-quantitative information on host abundance and may offer limited taxonomic resolution due to sequencing read length constraints ^3^. Fluorescence-activated cell sorting (FACS)-based approaches can be used to investigate gene transfer events but typically require genetically engineered reporter systems and are therefore not readily applicable to natural microbial communities ^14–16^. Consequently, a major methodological gap remains in the direct visualization of ARGs and host identity within individual cells in complex microbial communities.

GeneFISH had addressed some of these challenges by combining catalyzed reporter deposition – fluorescence *in situ* hybridization (CARD-FISH) for single-cell taxonomic identification using rRNA-targeted oligonucleotide probes with the detection of functional genes using digoxigenin-labelled ploynucleotide probes and anti-digoxigenin-horseradish peroxidase-mediated signal amplification ^17,18^. This approach has been successfully applied to diverse environmental microorganisms, including ammonia-oxidizing archaea in seawater ^18^ and denitrifying prokaryotes in groundwater ^19^, demonstrating its ability to directly associate functional genes with specific microbial taxa *in situ*. However, the multiple labeling, antibody detection and signal-amplification steps make the protocol labor-intensive and time-consuming. Direct-geneFISH was subsequently developed to simplify the procedure by directly coupling fluorophores to rRNA- and gene-targeting probes, reducing the protocol duration from more than 30 h to less than 6 h and enabling direct visualization, localization, and quantification of target genes within individual cells ^20,21^. The approach has been used to investigate functional diversity in marine planktonic bacteria ^22^, and microbial symbionts ^23^. Nevertheless, directly labelled rRNA probes may provide insufficient signal for robust host identification in environmental cells with low ribosome content. Thus, existing geneFISH approaches present a methodological trade-off between amplified, sensitive taxonomic identification and direct, quantitative gene detection.

Here, we address this trade-off by introducing combined-FISH, a sequential workflow that couples CARD-FISH amplification of rRNA signals for sensitive host identification with fluorophore-labelled polynucleotide probes for quantitative ARG detection. This combination enables ARG copies to be visualized and enumerated within taxonomically identified individual cells. As a model ARG, we selected the sulfonamide-resistant dihydropteroate synthase-encoding *sul*1, one of the most widely used indicators of anthropogenic AMR contamination in environmental compartments ^24,25^. We designed a set of polynucleotide probes targeting *sul*1 and established hybridization conditions compatible with joint 16S rRNA and ARG detection and quantification in the same cells. The method was validated in model strains carrying plasmid-borne *sul*1, where it enabled quantitative determination of gene copy numbers within individual host cells, applied to monitor plasmid-mediated gene transfer, and further extended to environmental biofilm communities to visualize *sul*1-carrying bacteria in complex microbial systems.

## Materials and Methods

### Model strains and plasmids

The following bacterial strains were used, all obtained from the German Collection of Microorganisms and Cell Cultures GmbH (DSMZ): *Acinetobacter defluvii* strain WCHA30 (DSM 107594) harboring the *sul*1 gene on the plasmid pOXA58_010030 (plasmid size: 355 kb; predicted mobility according to Plasmid Database ^26^: mobilizable; replicon type: not defined; relaxase type: MOB_P_; GenBank accession number: CP029396.2), *Escherichia coli* K12 J53 (DSM 5189) carrying the *sul*1 gene on the well-characterized plasmid R388 ^27,28^ (size: 33 kb; mobility: conjugative; replicon type: IncW; relaxase type: MOB_F_; GenBank accession: [NC_028464.1](https://www.ncbi.nlm.nih.gov/nuccore/CP029396.2?report=genbank)), and *Pseudomonas alloputida* KT2440 (DSM 6125, previously named *P. putida* KT2440). For brevity, the strain designations are not mentioned below.

Standard cultivation procedures were followed with *E. coli* grown in lysogeny broth (LB) or agar, *P. alloputida* grown in LB or DSM medium 1 (5.0 g peptone and 3.0 g meat extract per liter of distilled water at pH 7.0), and *A. defluvii* cultivated in DSM medium 1. For the two *sul*1-containing species, *A. defluvii* and *E. coli*, 100 µg/mL sulfadiazine (SDZ) was supplemented to the medium to maintain plasmid stability.

For combined-FISH, cells from mid-log phase (OD_600_ ≈ 0.5) were harvested by centrifugation at 10,000 rpm for 15 min (Centrifuge 5424/5424R, Eppendorf) and fixed by resuspension in 4% paraformaldehyde (PFA) in 1 × phosphate-buffered saline (PBS) for 1 hour at room temperature. Following fixation, cells were washed twice with 1× PBS, resuspended in 1 × PBS containing 50% ethanol, and stored at -20°C until further use.

### Conjugation between *E. coli* and *P. alloputida*

To test the applicability of combined-FISH for tracking conjugation and then to determine plasmid R388 copy numbers in *P. alloputida* transconjugants, conjugation assays between *E. coli* as the donor and *P. alloputida* as the recipient were carried out. SDZ was included in assays as a selection marker, with growth inhibition of *P. alloputida* at ≥16 µg SDZ /mL (Fig. S1).

Donor and recipient strains were grown individually in LB to an OD_600_ of approximately 0.5 each. Cells were harvested by centrifugation, washed twice with sterile LB, and resuspended in fresh LB. Donor and recipient suspensions were mixed with a cell ratio of approximately 1:1 (total volume of 15 mL LB per condition). Cultures were incubated statically at 30°C for 2 h ^29^. Then, cells were fixed with 4% PFA and stored at -20°C until analysis with combined-FISH. Conjugation efficiency was calculated as the number of observed transconjugant cells divided by the number of recipient cells.

To facilitate the generation of *P. alloputida* transconjugants for analysis of plasmid copy numbers in this bacterium, conjugation assays in LB were also carried out in which donor and recipient suspensions were added to 100 mL LB to achieve an initial OD_600_ of approximately 0.02. Cultures were incubated statically at 30°C for 12 h. Subsequently, aliquots were plated on cetrimide agar supplemented with 100 µg/mL SDZ and incubated at 30°C for 24 h to select for transconjugants (*Note*: the *qacEΔ1* gene on plasmid R388 confers tolerance to cetrimide in *E. coli* ^30^; however, the 100 µg cetrimide/mL used here is higher than the reported minimum inhibitory concentration of 64 µg/mL). Resulting colonies were purified by two successive transfers on fresh cetrimide agar with 100 µg/mL SDZ. Then, individual colonies were randomly selected, inoculated into fresh LB, and grown to an OD_600_ of approximately 0.5 before being harvested by centrifugation and fixation with PFA.

### Biofilm sample collection

Biofilm samples were collected from the discharge point of the Wernigerode-Silstedt Wastewater Treatment Plant (WWTP), Holtemme River, Germany (N51°51′59.4″, E10°51′43.3″), which was previously investigated for anthropogenic impact, including AMR, on small river systems ^31–33,34^. Riverbed stones containing microbial biofilms were collected and immersed in 50 mL filtered river water to maintain hydration during transport (4–5 stones per sterilized 250 mL glass bottle; filtered water was obtained by sequential filtration through a 0.45 µm filter and a 0.22 µm sterile syringe filter). All samples were transported to the laboratory within 3 hours in a thermostable container with cold packs.

Upon arrival at the laboratory, riverbed stones containing biofilm samples were incubated in parallel on a shaker at 20 rpm at room temperature for up to 48 hours. A control-subset of samples was spiked with *E. coli* with R388 at a final concentration of 8.4 × 10^7^ cells/mL. The spiking suspension was prepared from log phase *E. coli* (OD_600_ approximately 0.5) grown in LB supplemented with 100 µg/mL SDZ, standard harvesting by centrifugation, washing with 1× PBS, and resuspension in 1× PBS. To determine the cell density of the spiking suspension, 1 mL was fixed with 4% PFA for 4′,6-diamidino-2-phenylindole (DAPI) staining and enumerated using microscopy. From individual biofilm samples, cells were harvested at T0 (5–10 minutes after *E. coli* addition to the control sub-set), T24 (following 24 hours of incubation), and T48 (48 hours of incubation). Following incubations, the biofilm was collected by scraping of the stone surfaces using a sterile plastic inoculation loop and chemically fixed with 2% PFA prepared in 1× PBS, for 1h at room temperature. To remove the fixative, the samples were then transferred to sterile centrifuge tubes, centrifuged at 10,000 rpm for 15 minutes at 4°C, and washed twice with 1× PBS. The washed biofilm pellets were resuspended in 2 mL of 50% ethanol in 1× PBS, ultrasonicated in water bath (SONOREX DIGITEC DT 1028 CH, BANDELIN, Germany; 35 kHz; nominal ultrasonic power, 300 W) for three 20-min periods to disperse the cell, and stored at -20°C until further analysis.

### Probe design and fluorescent labeling of probes

#### rRNA probes

Standard rRNA oligonucleotide probes used in this study were: EUB338Ⅰ-Ⅲ (targeting almost all bacteria), Gam42a (targeting the 23S rRNA of *Gamaproteobacteria*), Gam42 comp (served as a competitive probe of Gam42a), PSE1284 (targeting the 16S rRNA of *Pseudomonas spp.*) and NONEUB (a negative probe for all rRNA probes), with sequences and formamide concentration used in hybridization shown in Table S1. The probes were covalently linked to horseradish peroxidase by Biomers.net GmbH (Ulm, Germany). Probes were reconstituted in ultrapure water to a stock concentration of 50 pmol/µL according to the manufacturer’s instructions and stored at -20°C.

#### Gene probes

For probe design, we downloaded all 23 *sul*1 reference sequences, which are complete coding sequence (CDS) and, if available, adjacent up- and downstream sequences, from the NCBI RefSeq database, and the *sul*1 reference sequence from the Comprehensive Antibiotic Resistance Database (CARD) ^35^. The *sul*1 sequences previously recovered from the Holtemme river ^33^ are identical to the CARD sequence and were therefore inherently included in probe design. All sequences were aligned using Geneious Prime software (https://www.geneious.com/) to identify the most conserved region without gaps and evenly distributed mismatches. PolyPro software was used to design probes targeting all aligned sequences within a 5% mismatch tolerance ^36^. Melting profiles of probes were predicted using MeltSim 1.0 to ensure compatibility for simultaneous hybridization ^37^. As negative control, a predicted methyltransferase gene from the archaeon *Acidilobus saccharovorans* 345-15 (GenBank accession CP001742.1) was chosen. Double-stranded DNA probes were synthesized by Integrated DNA Technologies (IDT, https://eu.idtdna.com/) and supplied as gBlocks Gene Fragments in dried powder form.

#### Probe Coverage Analysis

To assess probe coverage, accession numbers of 80 non-redundant Sul1 amino acid sequences were obtained from the structured ARG reference database (SARG) ^38^, followed by downloading 1552 corresponding CDS from NCBI’s GenBank. The hosts of these sequences were mainly *Escherichia* (25.8%), *Klebsiella* (23.2%), *Pseudomonas* (11.7%), *Acinetobacter* (8.6%), *Enterobacter* (7.2%), *Proteus* (6.1%), *Citrobacter* (4.0%) and *Salmonella* (3.4%) (see Fig. S2). Duplicate sequences were removed using BBMap ^39^ within Geneious Prime (*k*-mer seed length = 31). Dereplicated sequences were then aligned with the probe sequences in Geneious Prime, and the resulting matched regions were imported into PolyPro to calculate mismatch rates.

### Fluorescent labeling

To enable gene copy number enumeration in single cells, gene probes were labeled with the fluorescent dye Alexa Fluor 594 following a modified protocol based of Barrero-Canosa and Moraru (2021) ^21^. Individual gene probes were resuspended in TE buffer (1 mM EDTA, 5 mM Tris, pH 8.0) to a final concentration of 100 ng/µL, as recommended by the manufacturer. Labeling was performed using the Ulysis Alexa Fluor 594 Nucleic Acid Labeling Kit (Thermo Fisher Scientific, USA). We modified the standard protocol by using 1 µg of each probe per reaction, requiring 15 µL of labeling reagent (since the manufacturer does not provide dye concentration, we refer here to volumes of dye reagent). The choice of Alexa Fluor 594 was based on its strong and stable fluorescence signal and high detection efficiency ^20^. Labelled probes were purified using RNase-free Micro Bio-Spin 30 columns (Bio-Rad) according to the manufacturer’s instructions. Prior to purification, the original buffer in the column was exchanged to TE buffer according to the product specification. The base:dye ratio was calculated according to Barrero-Canosa and Moraru (2021) ^21^, giving 1 dye to every 22 bases for *sul*1 probe, and 1 dye to every 20 bases for the Nonsul probe.

### Combined rRNA CARD-FISH and direct-geneFISH protocol

The direct-geneFISH protocol was performed essentially as previously described ^21^, except that the mRNA hybridization step was omitted. The combined-FISH protocol consisted of the following steps: (1) Permeabilization, (2) Inactivation of endogenous peroxidases, (3) rRNA hybridization, (4) Tyramide signal amplification, (5) Washing, (6) Denaturation and gene hybridization, (7) Washing, and (8) Embedding and counterstaining with DAPI. A workflow schematic is provided in Fig. S3. Steps 2-5 were carried out according to the CARD-FISH protocol detailed in ^33^, while steps 6-8 were carried out as previously outlined for direct-geneFISH ^21^. Permeabilization varied on sample type: model organisms were treated with 0.5 mg/mL lysozyme, while biofilm samples were permeabilized with 10 mg/mL lysozyme and 60 U/mL achromopeptidase. Each gene probe was used at a concentration of 60 pg/µL, and the concentration of the Nonsul probe was 180 pg/µL. All FISH procedures were conducted on Superfrost^TM^Plus Adhesion Microscope Slides. And, all samples were mounted by ProLong™ Gold Antifade Mountant with DNA Stain DAPI (Thermo Fisher Scirntific, USA) before microscopy.

### Data collection and analysis

#### Microscopy

Following combined-FISH hybridization, samples were evaluated using a Zeiss Axio Observer 7 fluorescence microscope equipped with a Colibri.2 LED light source. DAPI and fluorophore-labeled probes were visualized using the following filter sets: DAPI (excitation 370 ± 40 nm, emission 425 nm, beam splitter 395 nm; filter set 62 HE) and Alexa Fluor 488 (excitation 474 ± 28 nm, emission 527 nm, beam splitter 495 nm; filter set 62 HE). Alexa Fluor 594 was visualized using filter set 64 HE (excitation 587 ± 25 nm, emission 647 ± 70 nm, beam splitter 605 nm).

Images were acquired with an Axiocam 503 mono black and white camera. To optimize image quality and signal-to-noise ratio, the following post-processing parameters were applied during acquisition: a noise filter (strength 3.0), and an unsharp mask (strength 3.0, radius 5.0). Camera gain was set to 3× to enhance sensitivity and minimize photobleaching. Exposure times were optimized for each fluorophore: DAPI, 50 ms; Alexa Fluor 488, 300-400 ms; and Alexa Fluor 594, 1000-4000 ms. A range of exposure times were tested, and images were selected based on optimal signal clarity and minimal background. Black and white images were pseudo-colored using ZEN 3.5 (blue edition) software (Zeiss).

### Image analysis & data interpretation

To optimize visualization of gene probe signals, histogram contrast and brightness were adjusted uniformly across all images. Thresholding parameters were consistent between experimental and negative control samples to ensure comparable assessment of potential non-specific signal. Gene detection efficiency was quantified using CellProfiler and ImageJ software on 5-10 images per condition, representing a minimum of 900 cells per culture. Detection efficiency was calculated as [(number of cells with gene signal / total cells analyzed) × 100] and expressed as the mean ± standard deviation across all fields of view (FoV).

To quantify gene copy number per cell, approximately 300-600 cells exhibiting strong signals in both rRNA and gene probe channels were manually counted using ZEN 3.5 software. Regions of interest (ROIs) were defined as circular selections around individual cells based on the rRNA signal.

Within each cell ROI, individual Sul1_3R signals were identified based on their punctate morphology and fluorescence intensity exceeding background levels and were manually delineated using the Draw Spline Contour tool in ZEN 3.5 software. Following the approach established for direct-geneFISH ^20^, individual spatially resolved Sul1_3R foci were considered proxies for single copies of the target gene and for plasmid copies.

### Statistical analysis

Statistical significance was determined using a two-tailed Student’s *t*-test, performed in Excel, to compare labeling efficiencies. Equal variances were confirmed using an F-test. Bar charts were generated using GraphPad Prism 10.

## Results

### Gene probe design, specificity, and coverage

For probe design, a total of 24 *sul*1 reference sequences from the curated, non-redundant NCBI RefSeq and CARD collections were used as template. The *sul*1 sequences previously recovered from the WWTP downstream site of the Holtemme River were identical to the CARD sequence ^33^. Based on the alignment of these sequences, an 819-bp highly conserved region (99.5% pairwise sequence identity) was selected for probe design. This region was subsequently divided into three consecutive 273-bp segments (regions 1–3), each serving as the template for designing one polynucleotide probe (Fig. S4 a). Sequences targeting all aligned *sul*1 genes within 5% mismatches and 5°C melting temperature (T_m_) difference were selected as candidate probes. Five candidate probes were identified for regions 1 and 2, and four were found for region 3 (Fig. S4 b). To enable simultaneous hybridization, the melting profile of candidate probes from each region was calculated and one probe was selected from each region such that the three selected probes exhibited closely matched melting temperatures. The probes were combined to form the final polynucleotide probe mix, Sul1_3R. (Fig. S4 c and Table S2). The G + C content of the three Sul1_3R probes ranged from 60.4 to 64.1%.

A 302 bp-long fragment of an archaeal methyltransferase gene was chosen as negative control probe, named Nonsul. This fragment has a similar GC content (60.3%) to the Sul1_3R probe set, which allows the same hybridization temperature as the Sul1_3R probe set. No match of the Nonsul polynucleotide probe sequence was found in a Blastn search in GenBank with “Bacteria” as query restriction.

To assess the coverage of the designed Sul1_3R probe set, we aligned each probe individually to 388 unique *sul*1 nucleotide sequences obtained from GenBank. The hosts of these sequences were mainly members of the *Enterobacteriaceae* (25.8%), *Pseudomonas spp*. (23.2%), and *Acinetobacter spp*. (11.7%). The alignments and mismatch calculation in PolyPro showed that the probes Sul1-region1, Sul1-region2, and Sul1-region3 target 98.2%, 99.7%, and 97.9% of the non-redundant (dereplicated) *sul*1 sequences, respectively (Fig. S5). Within a 5% mismatch threshold, the combination of the three probes in the hybridization mixture covers all unique *sul*1 nucleotide sequences in the NCBI RefSeq and CARD collections.

### Establishment of the combined-FISH protocol with model strains

To evaluate the applicability of the combined-FISH protocol across different bacterial hosts, we tested the Sul1_3R probe mix using two model strains carrying *sul*1 on genetically distinct plasmids: *E. coli* harboring the conjugative plasmid R388 and *A. defluvii* carrying the multidrug resistance plasmid pOXA58_010030. *P. alloputida*, which lacks *sul*1, served as a negative control. Taxonomic identification was performed using the Gam42a probe targeting the 23S rRNA of *Gammaproteobacteria*, while the unlabeled Bet42a probe was included as a competitor to assess hybridization specificity under the selected stringency conditions, which are relevant for subsequent applications to environmental samples. The combined-FISH workflow used to simultaneously detect *sul*1 and determine host identity is summarized in Fig. S3. Hybridization conditions were optimized according to established direct-geneFISH guidelines ^21^ while preserving cell integrity and morphology. Gene hybridization was performed using 55% formamide, followed by sequential washing under conditions appropriate for gene and rRNA probes.

Combined-FISH efficiently detected *sul*1 in both model strains, with signals observed in 82 ± 8.2% of *E. coli* cells and 84 ± 4.5% of *A. defluvii* cells (Fig. 1a and d). Signal intensities were predominantly between 4,000 and 8,000 arbitrary units (AU), while most Sul1_3R signals occupied an area of 0.1–0.3 µm² (Fig. 2a and b). Hybridization with the Nonsul probe resulted in low false-positive rates (6.0 ± 3.9% for *E. coli* and 1.0 ± 1.1% for *A. defluvii*), comparable to those previously reported for other genes by direct-geneFISH ^20,22^. No signal above background was detected in *P. alloputida* with either the Sul1_3R or Nonsul probe, confirming the specificity of the probe set (Fig. 1g).

**Fig 1.**
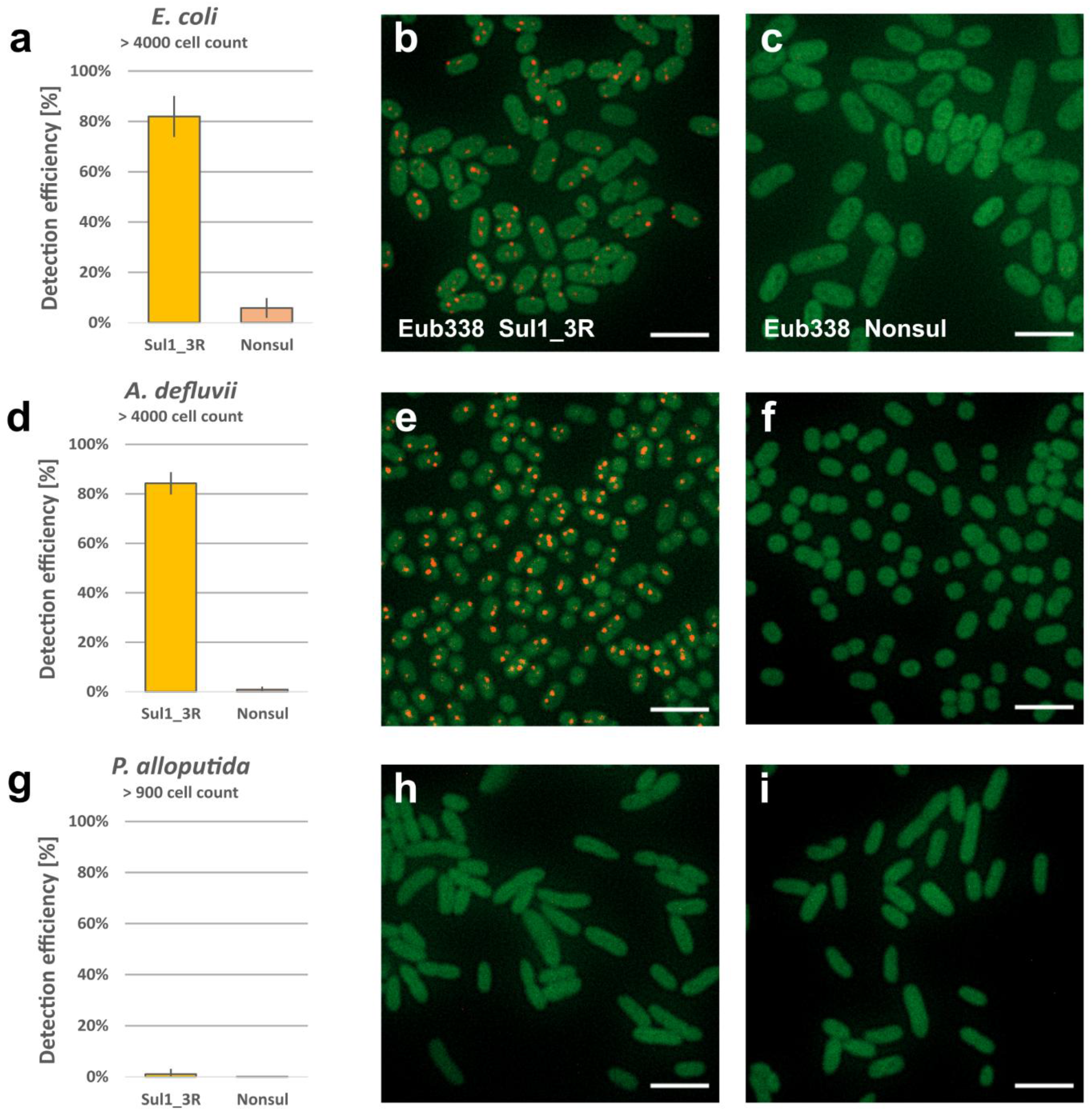
GeneFISH on model strains. a), d) and g) Detection efficiency of Sul1_3R and Nonsul probe on *E. coli*, *A. defluvii* and *P. alloputida*; b), e) and h) Overlay of rRNA and Sul1_3R signal of *E. coli*, *A. defluvii* and *P. alloputida*; respectively c), f) and i) Overlay of rRNA and Nonsul signal of *E. coli*, *A. defluvii* and *P. alloputida*, respectively. Scale bar: 5 µm

**Fig 2.**
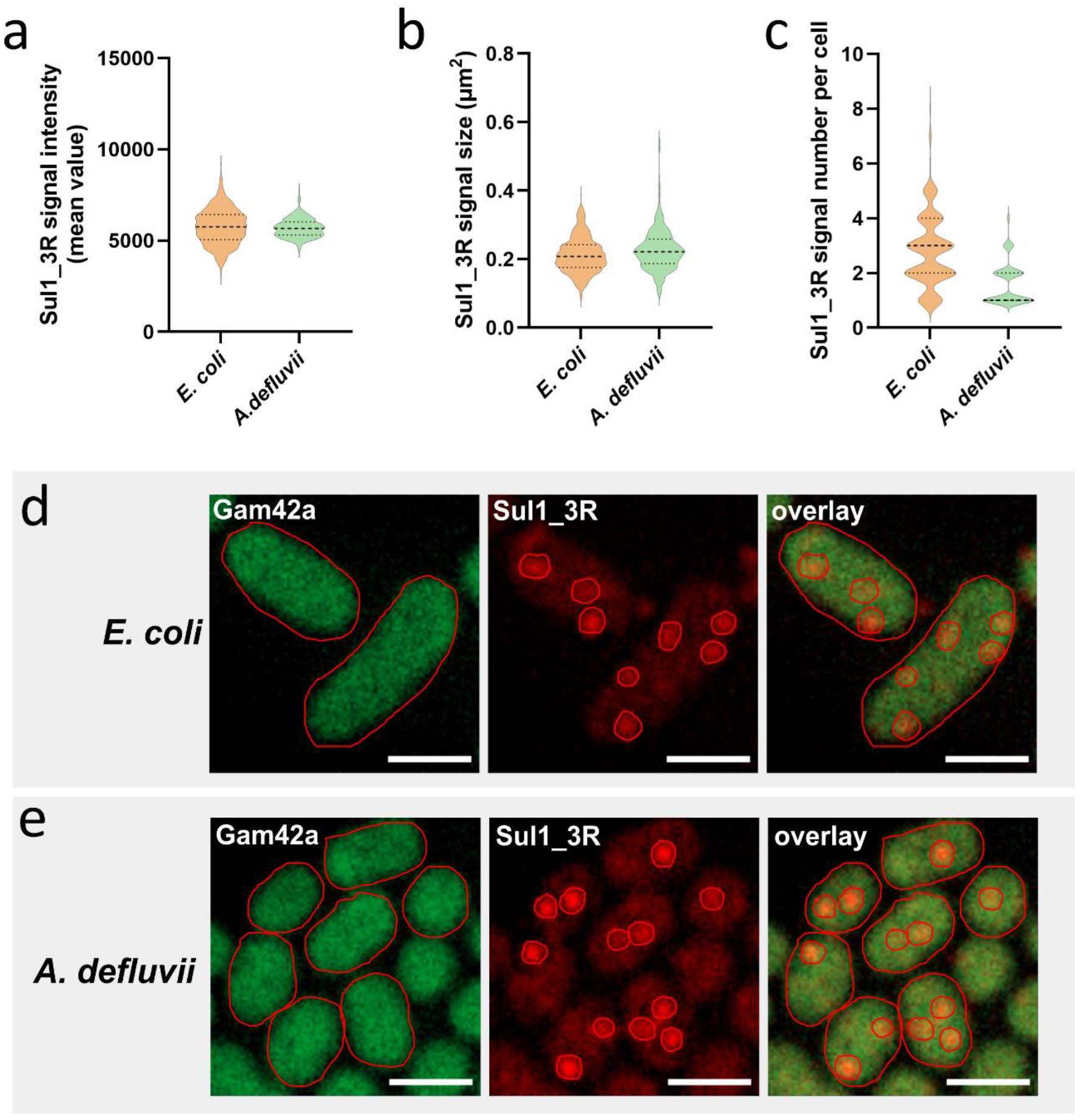
Sul1_3R probe signal per cell, signal size, and fluorescence intensity. a) Sul1_3R signal intensities in *E. coli* and *A. defluvii*; b) Sul1_3R signal size in *E. coli* and *A. defluvii*; c) Sul1_3R signal number in *E. coli* and *A. defluvii*; d) Example of quantifying the number and size of Sul1_3R signals (red) in *E. coli* (green); e) Example of quantifying the number and size of Sul1_3R signals (red) in *A. defluvii* (green). Scale bar: 2 μm

Following the principles established for direct-geneFISH ^20^, distinct Sul1_3R signal areas were used as proxy for *sul*1 copy numbers. Per *E. coli* cell with signal, we detected 1-8 gene copies (Fig. 2c and d, e), with 89.4% of those cells exhibiting 1–4 copies and the remainder having five or more copies. The predominant proportion of cells exhibiting two copies (30.9%), followed by cells with three (28.3%) and four copies (15.7%). In *A. defluvii*, combined-FISH revealed 1–4 *sul*1 gene copies per cell, with the majority of cells (54.2%) with discernable signal containing one copy, followed by 32.6% cells containing two copies and 11% cells with four copies. A small percentage (2.3%) of those cells exhibited more than four copies (Fig. 2c). The applicability of combined-FISH for detecting *sul*1-carrying bacteria was further validated on additional pure (Fig. S6) as well as mixed cultures (Fig. S7) that were available in our laboratory.

Following the successful proof-of-concept experiments with these model strains, we applied the combined-FISH protocol to track the transfer of the *sul*1-harboring conjugative plasmid R388 from *E. coli* to *P. alloputida* after 2 h of mating period. To distinguish between donor and recipient cells, we employed the PSE1284 probe, which specifically targets the 16S rRNA of *Pseudomonas spp.*, whereas donor *E. coli* cells were identified by DAPI staining. Combined-FISH detected *sul*1 signals in 4.5% ± 1.3% of the counted *P. alloputida* cells, a frequency that was significantly higher than the false-positive rate determined using the Nonsul probe (1.8% ± 2.2%, p < 0.05) (Fig. 3).

**Fig 3.**
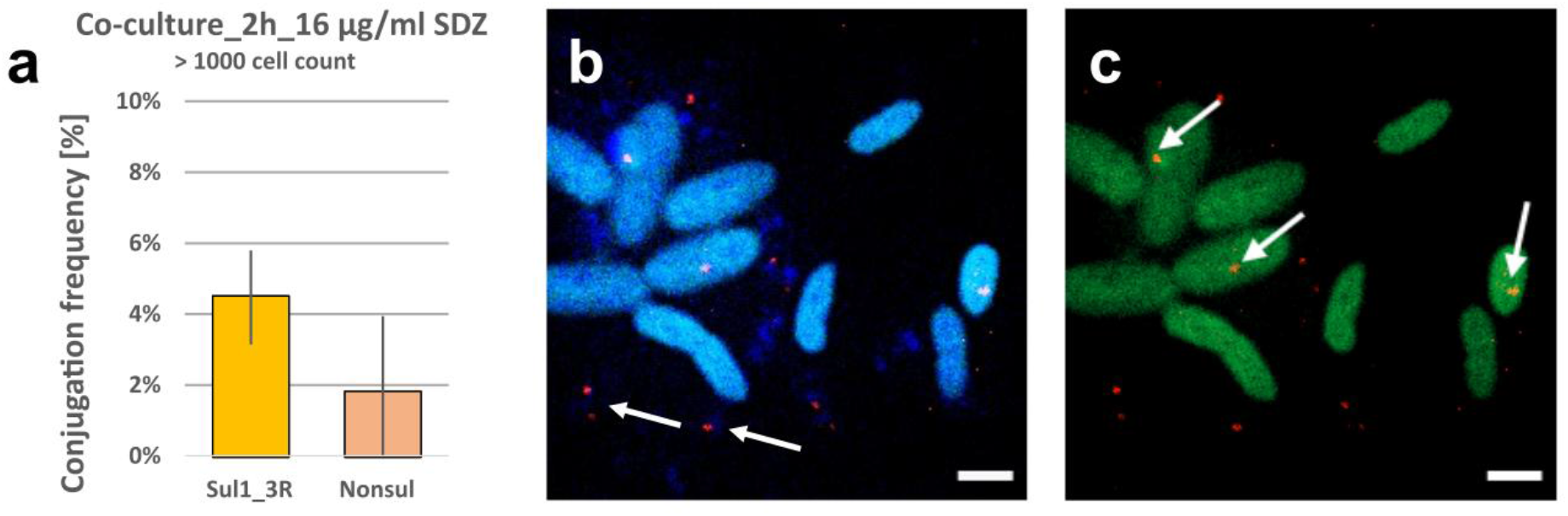
Combined-FISH after a 2-h conjugation assay with E. coli (DAPI staining) as donor of the R388 plasmid and *P. alloputida* (CARD-FISH staining of rRNA with probe PSE1284 and Alexa488 tyramides) as recipient. a) detection frequency; b) Overlay of DAPI, CARD-FISH (light blue) and Sul1_3R signals (red) of the co-culture; c) Overlay of CARD-FISH (green) and Sul1_3R signals (red) of the co-culture. The arrows in Fig. 3b point to *E. coli*, showing Sul1_3R signals (red) only with slight DAPI signals (due to strong denaturation) in their vicinity, while the arrows in Fig. 3c point to Sul1_3R signals (red dots) in apparent *P. alloputida* transconjugants (green). Scale bar: 2 μm.

To determine the copy number of the *sul*1-harboring R388 plasmid in *P. alloputida*, transconjugants obtained after overnight conjugation and antibiotic selection were analyzed by combined-FISH. Signals of *sul*1 were detected in 55% ± 7.1% of cells (Nonsul: 3.9% ± 1.6%) (Fig. 4a, b and c), ranging from 1 to 14 signals per transconjugant, 67% of which containing 2–6 signals (Fig. 4d), indicating a slightly higher plasmid copy number than that observed in *E. coli*.

**Fig 4.**
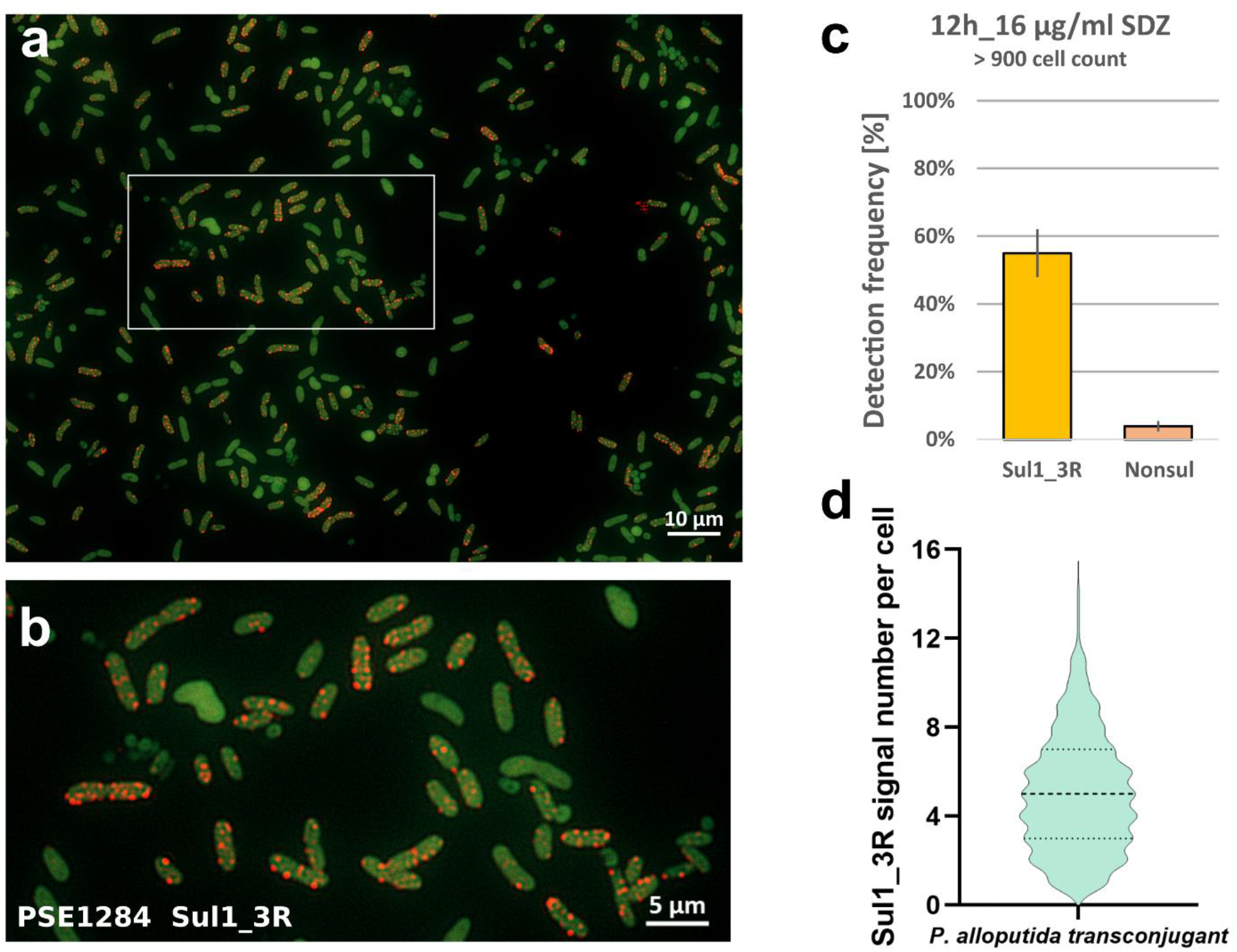
Combined-FISH on *P. alloputida* cells with R388 after successive transfer. a) overlay of rRNA (green) and Sul1_3R signal of cells (red), (whole field of view); b) overlay of rRNA and Sul1_3R signal of cells (cropped area); c) Detection frequency; d) Sul1_3R signal number per transconjugant cell.

### Detecting *sul*1-harboring bacteria in river biofilm

Next, we applied combined-FISH on biofilm samples from the Holtemme river, previously surveyed for AMR ^31,33,34^, to test the capability of the method for visualizing *sul*1-harboring bacteria in complex environmental samples. As positive controls, we used *E. coli* with R388 provided as axenic culture and added to river biofilm samples. The latter control was to test for potential inhibition by non-cellular biofilm material. The Gam42a probe was used for taxonomic identification of Gammaproteobacteria.

In axenic *E. coli* samples, detection efficiency was up to 94% (Fig. 5a and b), thereby better than in the proof-of-principle assays reported above. In biofilm samples inoculated with *E. coli*, the proportion of Gammaproteobacterial cells exhibiting a Sul1_3R signal was 57.5 ± 8.7% at T0 (Nonsul: 2.0 ± 1.6%), declined over 24 h to 30.4 ± 9.6% (T24; Nonsul: 8.4 ± 0.5%), and was at background levels (1.2 ± 1.1%) after 48 h (Nonsul: 2.7 ± 0.5%) (Fig. 5a). This control showed that detection in complex samples is feasible. Only a few cells displaying an *E. coli*-like morphology remained after 48 h of incubation (Fig. 5c), suggesting that the introduced *E. coli* cells with its *sul*1-harboring R388 plasmid did not persist in the river biofilm community under the conditions tested.

**Fig 5.**
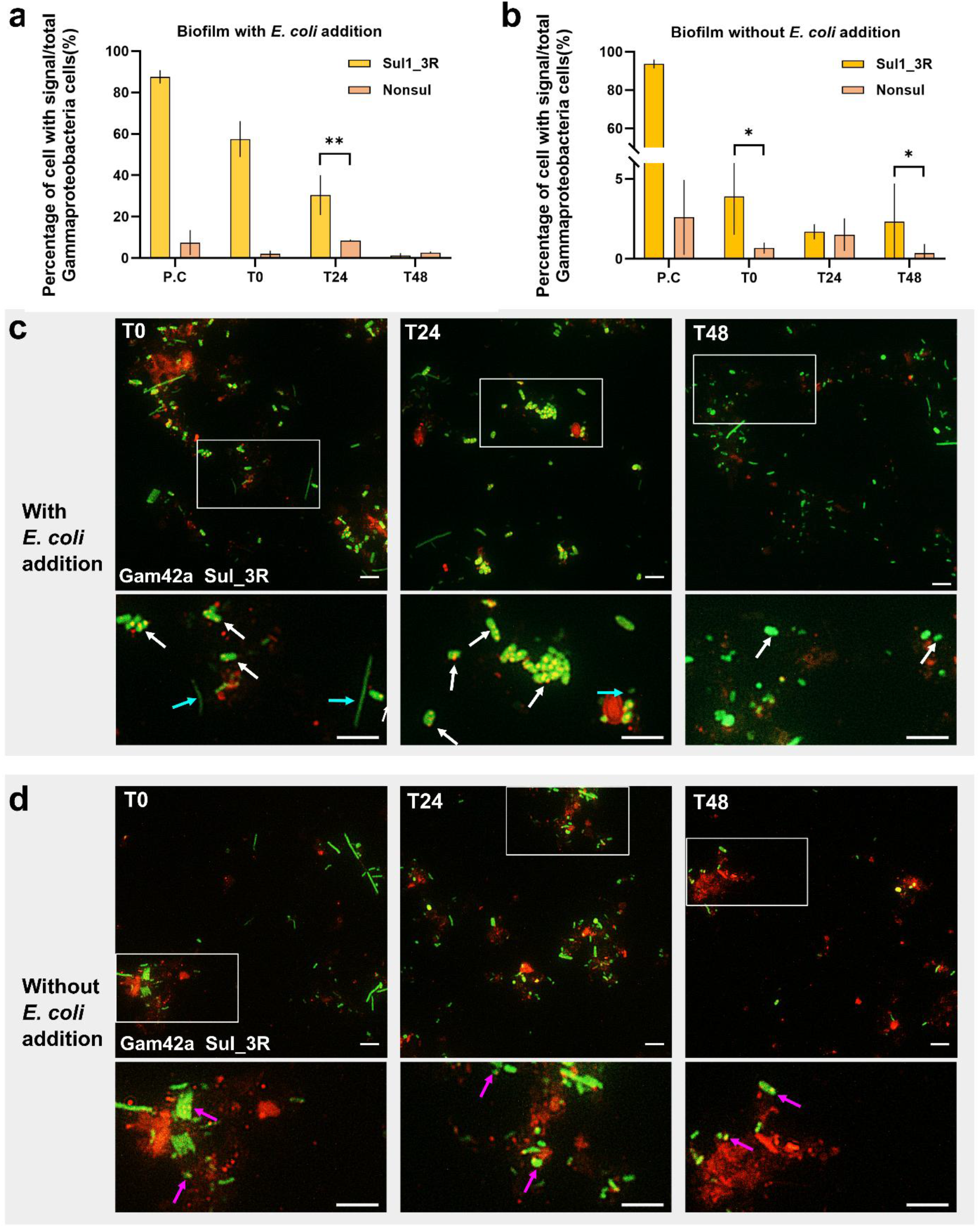
Fate of *E. coli* containing *sul1* in river biofilm. The percentage of cells with Sul_3R probe signal (dot like red) versus total Gamma-proteobacterial cells (green) in biofilm sample with *E. coli* addition (green) (a) and without E. coli (b); Representative fields of view (FoVs) of samples with *E. coli* addition (c) and without *E. coli* (d) at each sampling time point. P.C is the positive control that contains pure culture of *sul1* containing *E. coli*. White arrows pointing to the *E. coli* with Sul1_3R signal, blue arrows pointing to the *Gammaproteobacteria* inhabiting river biofilm, and pink arrows pointing to putative *E. coli* cells. Scale bar: 5 μm.

In the native river biofilm samples without *E. coli* addition, a proportion of 3.9 ± 2.4% of the Gammaproteobacterial cells were co-labelled with the Sul1_3R and Gam42a probes at T0 (Nonsul: 0.7 ± 0.3%; p < 0.05). At T24, the proportion of cells with Sul1_3R and Nonsul signal were about the same (1.7 ± 0.5 % vs. 1.5 ± 1.0 %), while there were statistically more cells labelled with Sul1_3R than with Nonsul (2.3 ± 2.4 % vs. 0.4 ± 0.6 %; p < 0.05) at T48 (Fig. 5b). Images of cells from the Holtemme river biofilm co-labeled with Sul1_3R and Gam42a probes are shown in Fig. 5d.

## Discussion

The combined-FISH protocol developed here integrates the quantitative capabilities of direct-geneFISH with CARD-FISH-amplified rRNA detection for sensitive host identification, thereby enabling simultaneous visualization of host identity and quantitative detection of ARG copy numbers at the single-cell level. By directly labeling polynucleotide probes with fluorophores, as in direct-geneFISH, the method permits the enumeration of individual gene copies within single cells while avoiding the tyramide signal amplification step required for gene detection in the original geneFISH protocol ^18,20^. This substantially reduces protocol complexity and processing time. A major challenge in combining rRNA and gene detection in a single hybridization protocol lies in the requirement for compatible hybridization stringencies, particularly with respect to formamide concentrations during hybridization and salt concentrations during washing. In our study, the relatively high G+C content of the Sul1_3R probes (60–64%) required hybridization conditions corresponding to approximately 55% formamide, which exceeded the optimal conditions for the rRNA-targeted probes employed here. Simultaneous hybridization under compromise conditions could therefore have reduced either rRNA labeling efficiency or gene specificity. To overcome this limitation, we adopted a sequential hybridization strategy similar to that used in the original geneFISH protocol, performing CARD-FISH and gene hybridization independently. This approach preserved the high specificity required for gene detection while maintaining the flexibility of CARD-FISH for taxonomic identification. Assuming that plasmid R388 with its *sul*1 gene was maintained by all *E. coli* cells (previously, 100% of cells retained the plasmid over 80 generations ^40^), we obtained hybridization efficiencies ranging from around 82% to up to 94% (positive control in Fig. 5b). This efficiency is similar to previous reports for single-copy gene detection in pure cultures using geneFISH and direct-geneFISH^18,20,22^, where a small fraction of cells were false negatives arising from incomplete probe accessibility or reduced hybridization efficiencies ^18,20^. The hybridization efficiencies in this study indicates that the additional CARD-FISH step does not substantially compromise gene detection performance.

Assuming that intracellular transposition of *sul*1 from plasmid to chromosome had not occurred in the model organisms, the combined-FISH results indicated heterogeneity in the plasmid copy number among *E. coli* cells (1 to 8 copies per cell, 59% of cells with 2 or 3 copies), *P. alloputida* after conjugation with plasmid R388 (1 to 14 copies, with 67% of cells containing 2–6 copies), and *A. defluvii* (1 to 4 copies, with 54.2% of cells containing 1 copy). These numbers are similar to previous estimates of the respective plasmid copy numbers in *E. coli* and *A. defluvii*. The average copy number of a *gfp*-labelled derivative of plasmid R388 in *E. coli* was estimated by fluorescence microscopy to be about 5, ranging from 1 to 10 depending on cell size ^40^, and the copy number of plasmid pOXA58_010030 in *A. defluvii* relative to chromosome copy number was estimated based on sequencing read coverage to be around one ^41^, fitting with the plasmid’s large size of 355 kb.

As with other approaches for estimating gene copy numbers, the interpretation of individual fluorescent foci as plasmid copies is subject to methodological limitations, including the spatial resolution of fluorescence microscopy and the potential overlap of adjacent signals, which may lead to an underestimation of abundance of gene and plasmid. Here, the low copy numbers fall within a range where individual fluorescent foci can generally be resolved using conventional fluorescence microscopy, particularly in rod-shaped bacterial cells ^20^. Accurate quantification would become increasingly challenging for high-copy-number plasmids, where the overlap of multiple signals within the same cell would prevent reliable discrimination of individual copies. Thus, the combined-FISH approach is particularly well suited for the investigation of low-copy-number plasmids and their dynamics at single-cell resolution. The method could be a valuable alternative or complement to existing methods for plasmid copy number determination ^42,43^, especially since combined-FISH does not rely on genetic modification of the plasmid for detection.

Traditional methods such as colony PCR for studying plasmid-mediated ARG transfer have culturing prerequisites, e.g. selective or at least differential agar specific to the recipient strain is typically required, and the donor strain may have to carry a plasmid containing ARGs that confer resistance to specific antibiotics ^44–46^. A recently published high-throughput method can analyze multiple conjugation events in 96-well plates ^47^. However, this method still depends on culturing. The requirements make these methods impractical or unsuitable for environmental samples, as they rely on specialized media and growth conditions that may not accurately reflect those of the natural habitats. Advanced methods like FACS and microfluidics are fluorescence-based techniques that require genetic modification of the donor strain ^14,15,48^. Compared to these methods, the combined-FISH protocol can track plasmid-borne ARG transfer events without multiple prerequisites. For low-frequency transfer events, distinguishing true positives from false positives remains challenging.

Furthermore, we applied combined-FISH in a time-course assay to native river biofilm samples from a WWTP discharge point and found up to approximately 4% of the biofilm-bacteria were co-labeled with Sul1_3R and Gam42a at two out of three time points. At the third, the Sul1_3R signal was not statistically different from the negative control with the Nonsul probe. The percentages, especially at time point T0, agree well with the previous determination of *sul*1 by qPCR at a relative abundance of around 3 × 10⁻² copies/16S rRNA gene ^34^. Furthermore, although the taxonomic *sul*1 host profile at the site is unknown, it seems reasonable to assume that the majority of the hosts were *Gammaproteobacteria* considering previous knowledge from sequenced genomes ^49^ and epicPCR analysis of wastewater ^11^. At the sampling site, this class constituted approximately 1/3 of all bacteria ^31^. It seems also reasonable to assume that the ratio of *sul*1 copies to 16S rRNA gene copies per cell at the site was around one. Given these enumerations and estimates, we suggest applying the combined-FISH protocol to environmental samples with relative abundances of the target gene exceeding 1 × 10⁻² copies per 16S rRNA gene, as determined by qPCR or metagenomics. Together, these findings establish combined-FISH as a powerful tool for quantitatively linking ARGs and host cells *in situ*, as well as for determining plasmid copy numbers. Future developments of this approach could expand its applicability in environmental and clinical antimicrobial resistance surveillance. Accurate assessment of ARG-associated risks requires direct identification of high-risk resistance determinants within their pathogenic hosts. Building on the combined-FISH framework, polynucleotide probes could be designed for selected ARGs classified as high-risk to human health, as recently proposed by Zhang et al. ^50^, enabling detection of multiple priority resistance genes under optimized hybridization conditions. The establishment of a curated ARG probe database would further facilitate the implementation of this approach across diverse microbial ecosystems. Combining ARG-targeting probes with pathogen-specific rRNA markers would enable direct in situ assessment of antibiotic resistance risks in environmentally and clinically relevant samples. Integration with flow cytometry could further increase throughput, while coupling with high-taxonomic-resolution approaches such as HiPR-FISH ^51^ would allow spatial localization of high-risk ARGs across multiple host populations within complex microbial communities. Overall, these developments position the combined-FISH approach as a versatile platform for quantitatively assessing ARG risks and linking resistance genes to microbial identity *in situ*.

## Acknowledgement

We would like to express our sincere gratitude to Jasmin Vogt for providing training in CARD-FISH, Chiththaka Chaturanga Bandara for valuable assistance with fluorescence imaging, and Kristin Lindstaedt and Sebastian Roether for their support with image data analysis. We also gratefully acknowledge the ProVIS platform for providing access to the research infrastructure used in this study.

