## Supplementary information for "Single-cell imaging links antibiotic resistance gene copy number and bacterial host identity"

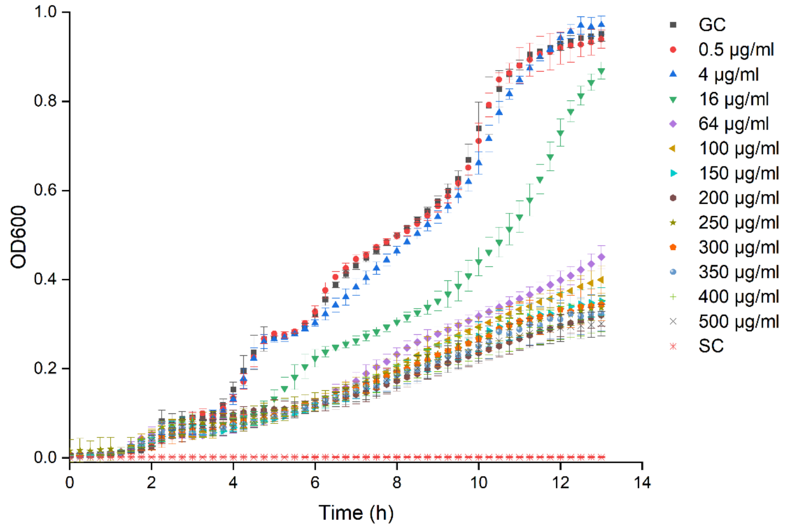

**Fig. S1** Minimum inhibitory concentration (MIC) test for *Pseudomonas alloputida* under different SDZ concentrations. GC: growth control (without SDZ addition); SC: sterile control (without bacteria addition)

**Table S1** List of rRNA probes used for combined-FISH

| **Target gene** | **Specificity** | **Sequence (5’–3’)** | **Size (bp)** | **Formamide concentration (%)** | **References** |
|---|---|---|---|---|---|
| Eub338I | 16S rRNA of most Bacteria | GCT GCC TCC CGT AGG AGT | 18 | 0-50 | [1] |
| Eub338II | 16S rRNA of *Planctomycetes* | GCA GCC ACC CGT AGG TGT | 18 | 0-50 | [2] |
| Eub338III | 16S rRNA of *Verrucomicrobium* | GCT GCC ACC CGT AGG TGT | 18 | 0-50 | [2] |
| EUB338I-III | Bacteria | Mixture of the probes EUB338, Eub338II and Eub338III | | 0-50 | [2] |
| Gam42a | 23S rRNA of *Gamaproteobacteria* | GCC TTC CCA CAT CGT TT | 17 | 35 | [3] |
| Gam42 comp | Competitive probe of Gam42a | GCC TTC CCA CTT CGT TT | 17 | 35 | [3,4] |
| PSE1284 | 16S rRNA of *Pseudomonas spp.* | GAT CCG GAC TAC GAT CGG TTT | 21 | 30 | [5] |
| NONEUB | Nonsense of EUB338 (The NON338 has the | ACT CCT ACG GGA GGC AGC | 18 | / | [6] |

| | |
|---|---|
| | same sequence and sense as the rRNA) |

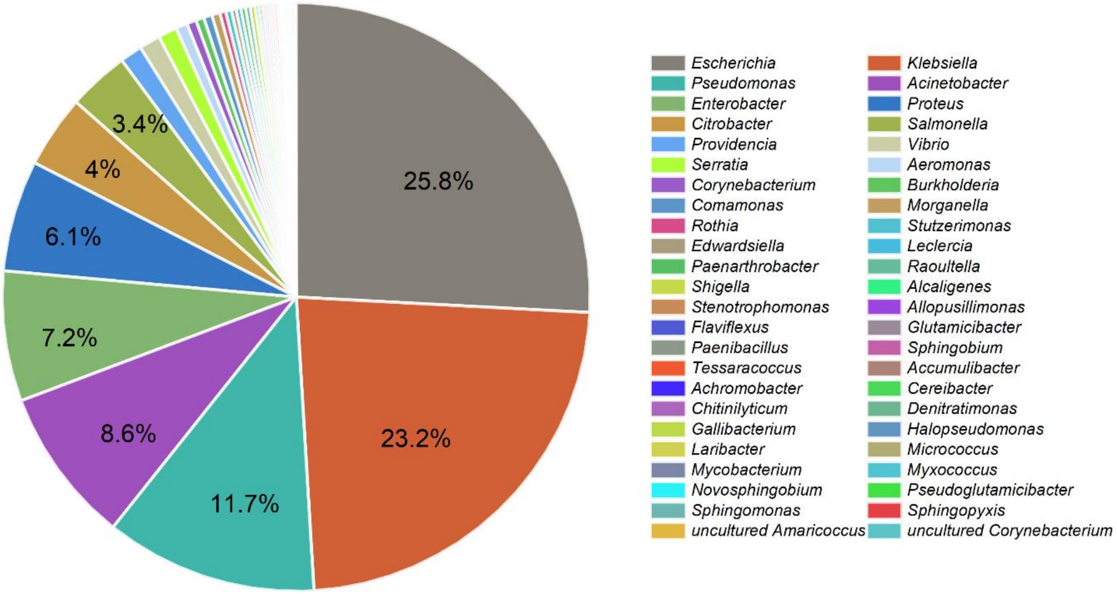

**Fig. S2** Host information and their proportion of 1552 *sul*1 sequences downloaded from NCBI according to *sul*1 reference collection from SARG database

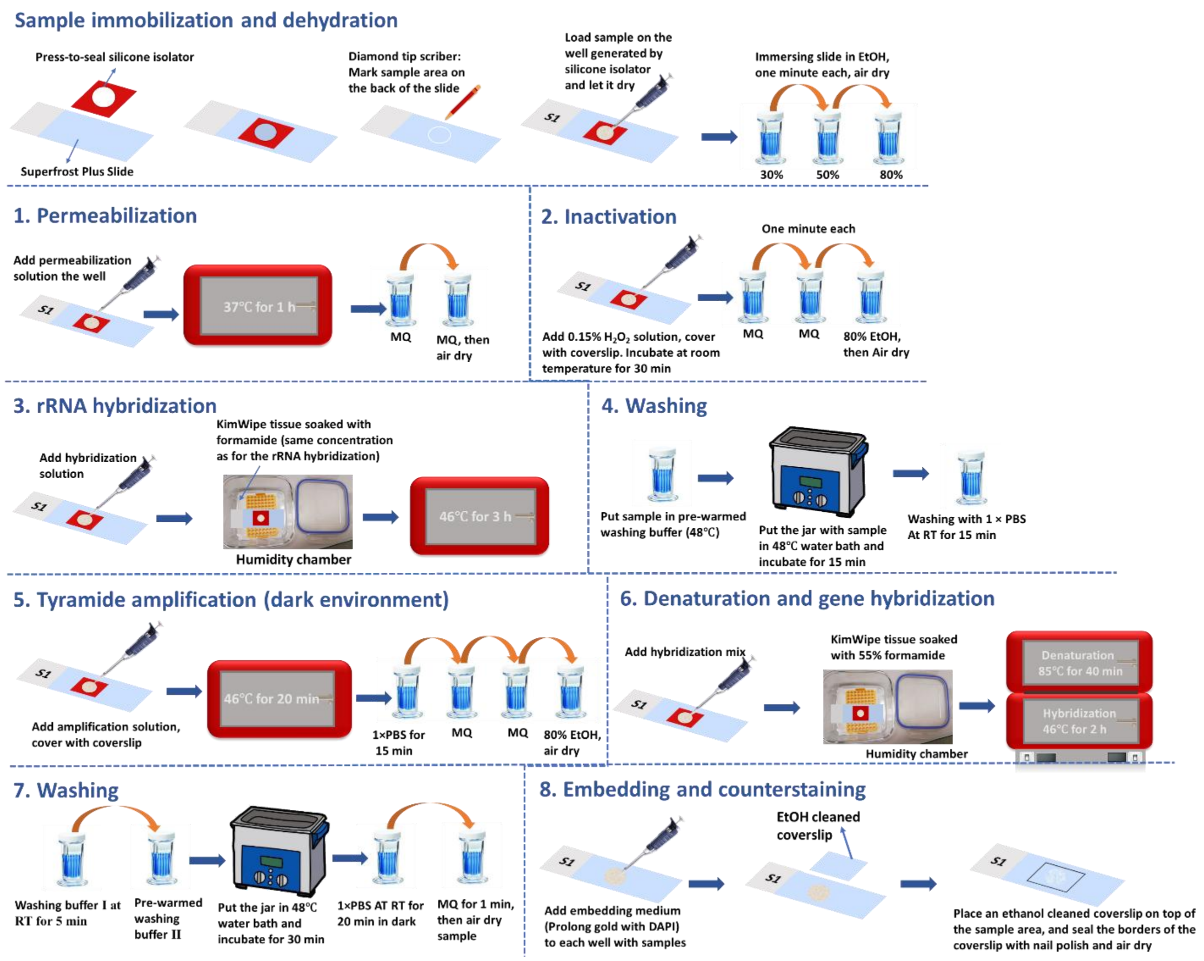

**Fig. S3** Detailed workflow of the combined-FISH protocol.

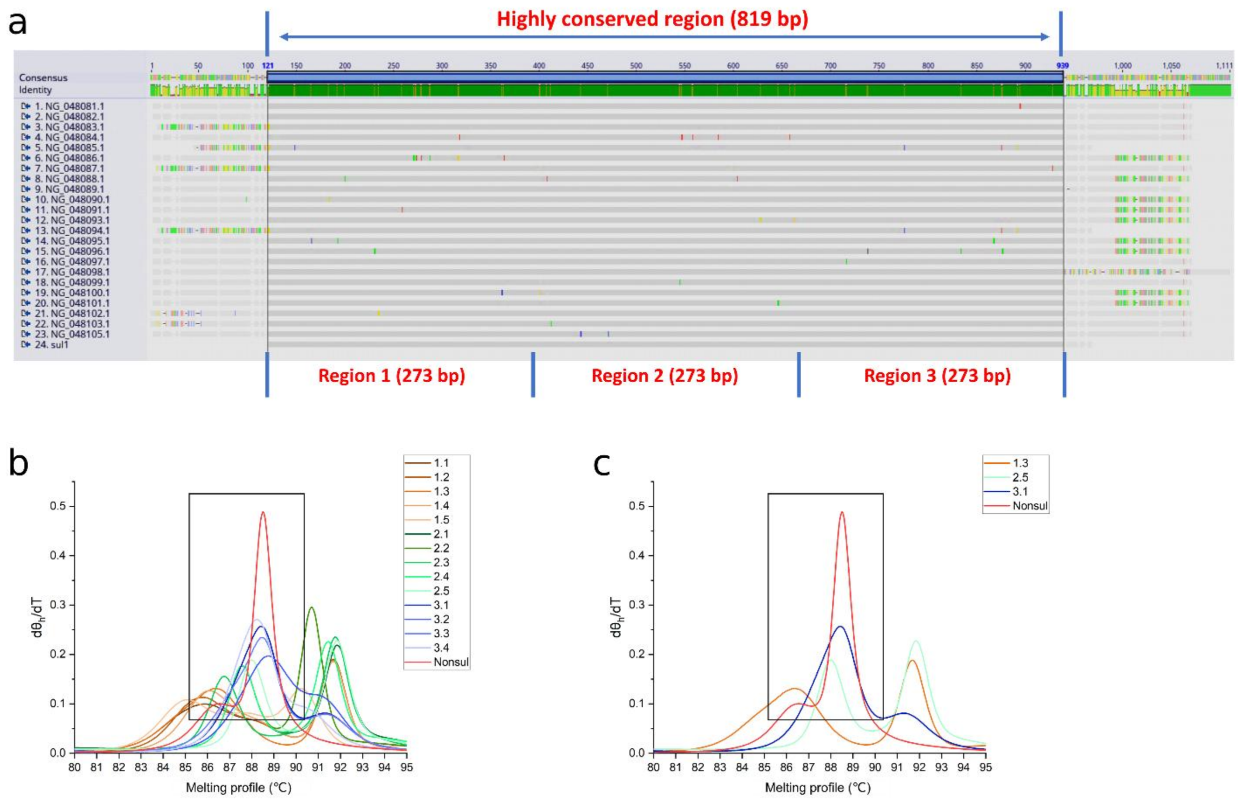

**Fig. S4** Alignment of sequences and melting profiles of probes. a) alignment of sequences of sulfonamide-resistant dihydropteroate synthase *sul1* gene family from NCBI (number 1 -23) and *sul1* reference sequence from CARD database (number 24). Note: In the green bar at the top, green signifies 100% identity across a column. Within the alignment itself, identical sequences are marked in grey, while differences are highlighted with colored lines; b) melting profiles of all candidate probes for region 1, 2 and 3; and c) melting profiles of final probes for region 1, 2 and 3. $\theta_h$ is the calculated fraction of base pairs remaining in the helical state. The former number in the legend represents the region.

**Table S2** Sequence information of designed probes

| Gene name | Sequence | G + C content (%) |
|---|---|---|
| Sul1-region1 | ATGGTGACGGTGTTCGGCATTCTGAATCTCACCGAGGACTCCTTCTTCGAT GAGAGCCGGCGGCTAGACCCCGCCGGCGCTGTCACCGCGGCGATCGAAAT GCTGCGAGTCGGATCAGACGTCGTGGATGTCGGACCGGCCGCCAGCCATC CGGACGCGAGGCCTGTATCGCCGGCCGATGAGATCAGACGTATTGCACCG CTCTTAGACGCCCTGTCCGATCAGATGCACCGTGTTTCAATCGACAGCTTC CAACCGGAAACCCAGCGCTAT | 61.2 |
| Sul1-region2 | GCGCTCAGGCGCGGCGTGGGCTACCTGAACGATATCCAAGGATTTCCTGA CCCTGCGCTCTATCCCGATATTGCTGAGGCGGACTGCAGGCTGGTGGTTAT GCACTCAGCGCAGCGGGATGGCATCGCCACCCGCACCGGTCACCTTCGAC CCGAAGACGCGCTCGACGAGATTGTGCGGTTCTTCGAGGCGCGGGTTTCC GCCTTGCGACGGAGCGGGGTCGCTGCCGACCGGCTCATCCTCGATCCGGG GATGGGATTTTTCTTGAGCCCC | 64.1 |
| Sul1-region3 | GCACCGGAAACATCGCTGCACGTGCTGTCGAACCTTCAAAAGCTGAAGTC GGCGTTGGGGCTTCCGCTATTGGTCTCGGTGTCGCGGAAATCCTTCTTGGG CGCCACCGTTGGCCTTCCTGTAAAGGATCTGGGTCCAGCGAGCCTTGCGGC GGAACTTCACGCGATCGGCAATGGCGCTGACTACGTCCGCACCCACGCGC CTGGAGATCTGCGAAGCGCAATCACCATCTCGGAAACCCTCGCGAAATTT CGCAGTCGCGACGCCAGAGAC | 60.4 |

| | | |
|---|---|---|
| Nonsul | GGTGGAGCTCAGGACGCCCACGTCATACTACAAGCTGGTGCCCTTGGAGG GCGGCGCTCCAACCCTGGAAATCAACGGCATACACATGCACAGGATCTCC GGCACCACGCCCCTAAGGGATGCCGAACTTAAGGTAAAGGCCGCTGGCGT GAGAAGAGGGGACAAGGTCCTCGAGATAGGCACGGGCTTAGGCTACACC GCCGCTGAGGCCCTCAAGGCGGGAGCTAAGGTCATCAGTGTGGAGCTCAG GGAAGAGGTGCTCTGGATTGCTGAGAGAAACCCGTGGAGCTTTCGGCTTA GGG | 60.3 |

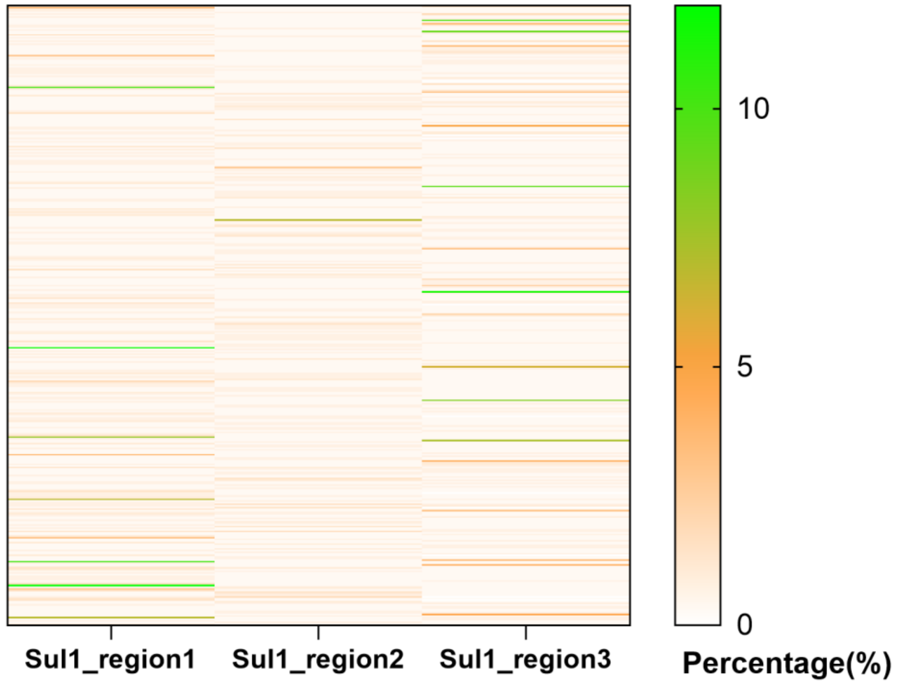

**Fig. S5** Mismatch percentage of each probe targeting each of the 388 non-redundant *sul1* nucleotide sequences.

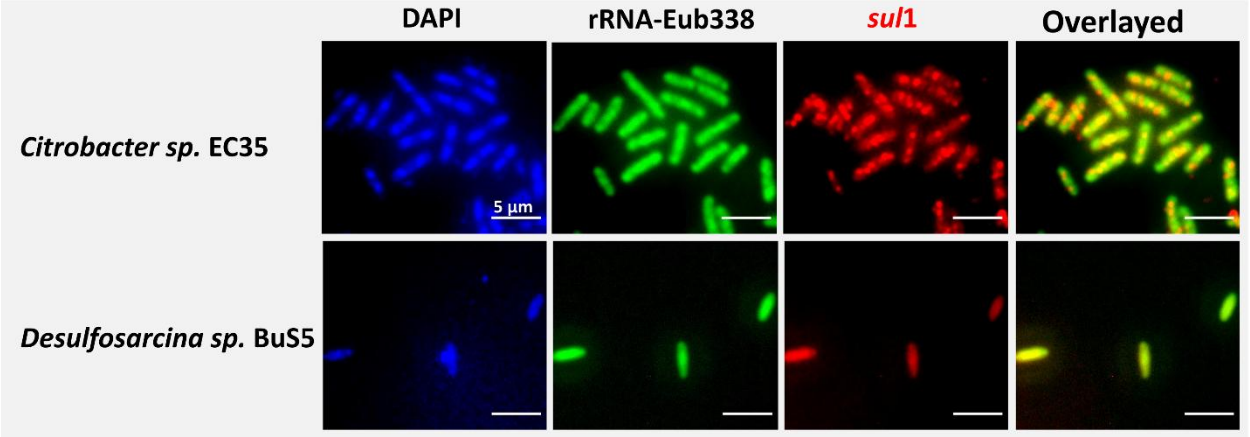

**Fig. S6** Additional model organisms used to evaluate the combined-FISH approach. *Citrobacter sp.* strain EC35 (obtained from the UFZ culture collection) that harbor the *sul*1 gene, whereas the *Desulfosarcina* related Butane oxidizing BuS 5 strain (from our laboratory), lacking *sul*1, served as the negative control.

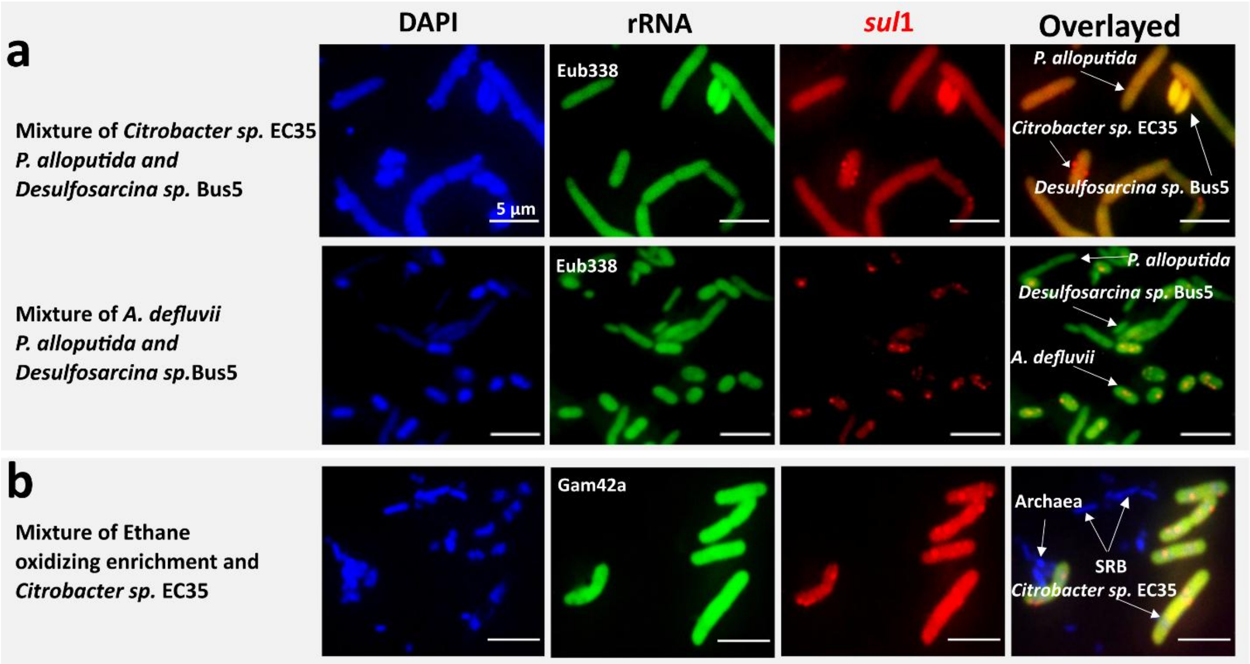

**Fig. S7** Various mixed combinations of the model/laboratory organisms with and without the *sul*1 gene hybridized with various 16S rRNA probes: Eub338 (a) and Gam42a (b). The Ethane-oxidizing enrichment culture consisted primarily of two phylotypes of *Desulfosarcina*-affiliated sulfate-reducing bacteria (SRB) and the archaeon "*Candidatus* Argoarchaeum ethanivorans" [7] that is from our laboratory. Strains of model/laboratory organisms and enrichment culture were grown separately, chemically fixed, and combined on filters during the filtration step and prior to the combined-FISH hybridizations.
